# Symbiont-mediated shifts in cuticular hydrocarbon profiles reduce female attractiveness after mating

**DOI:** 10.64898/2026.08.24.745886

**Authors:** Amir H. Tourani, Alihan Katlav, James M Cook, John Hunt, Sajjad Reyhani Haghighi, Shawan Karan, Markus Riegler

## Abstract

Males can influence future mating interactions of females after copulation by changing female signals that subsequent males will encounter. In insects, such effects commonly involve cuticular hydrocarbons (CHCs), but whether heritable microbial symbionts contribute to post-mating chemical signalling remains largely unknown. Kelly’s citrus thrips, *Pezothrips kellyanus*, provides an ideal system to address this question because reproductive compatibility is shaped by common arthropod endosymbionts. Across *P. kellyanus* populations, *Cardinium* occurs in almost all individuals whereas *Wolbachia* varies in prevalence and appears to spread by cytoplasmic incompatibility (CI). Yet, females with only *Cardinium* (C) avoid incompatible males carrying both *Cardinium* and *Wolbachia* (CW), and this discrimination is linked to the distinct CHC profile of CW males. Here, we tested whether this endosymbiont-associated male perfume persists on females beyond copulation by altering the female CHC profile and subsequent male mating behaviour. Using behavioural assays and GC–MS-based CHC profiling, we found that, independent of female endosymbiont association, females first mated with CW males received fewer antennal contacts and mating attempts from subsequent C males. Furthermore, mating remodelled female CHC profiles, while mating with CW males produced a distinctive post-mating chemical signature. Most notably, tridecane, previously detected only in CW males, occurred exclusively in females mated with CW males. Our findings show that endosymbionts can alter mated female CHCs and influence future sexual communication between male and female hosts. These findings reveal a previously unrecognised post-mating route through which endosymbionts reshape sexual communication, with potential consequences for reproductive compatibility and symbiont transmission dynamics.

## Introduction

For males, successful mating does not always guarantee paternity, because females often mate multiply and sperm acquired from different males may compete or be used differently. As a result, selection can favour male strategies that continue to influence female behaviour, signalling and reproductive interactions after copulation (Chapman *et al*., 2003, Ejima *et al*., 2007). Across animals, males have evolved diverse strategies to limit competition from future rivals after mating and maximise their own paternity. Examples are the transfer of seminal proteins during mating that suppress female receptivity to future matings (Sirot *et al*., 2011), mating plugs that interfere with the storage or use of sperm from subsequent mates (Devine, 1977; Saad *et al*., 2018) and/or changes in female signalling that reduce attractiveness to subsequent rival males (Thomas, 2011). Such post-mating effects are well documented across insects (Hollis *et al*., 2019), spiders (Schneider & Lesmono, 2009) and vertebrates (Vieites *et al*., 2004), and can arise from sexual conflict, where male paternity protection can oppose female control over remating (Parker, 1979; Arnqvist & Rowe, 2005).

Post-mating changes in female chemical signals can arise through different mechanisms, but they often have the same key effect - altering how later-arriving males perceive and respond to already mated females. By modifying the chemical cues associated with female identity, attractiveness or mating status, these changes can influence whether subsequent males detect already mated females, assess them as suitable mates and attempt further mating. Such chemically mediated reproductive responses occur across diverse animal taxa. For example, male courtship pheromones increase female receptivity in plethodontid salamanders (Houck & Reagan, 1990), female skin-lipid pheromones signal attractiveness and mating status in red-sided garter snakes (O’Donnell *et al*., 2004), and female vaginal secretions stimulate male sexual behaviour in hamsters (Macrides *et al*., 1984).

In insects, some of the most important chemical signalling compounds are cuticular hydrocarbons (CHCs), which function in both waterproofing of the cuticle and sexual communication (Howard and Blomquist, 2005; Blomquist & Bagnères, 2010). Because CHCs are displayed directly on the body surface, mating provides an opportunity for these chemical signatures to be physically altered through contact between males and females (Weddle *et al*., 2013). In this way, males may transfer components of their own CHC profile onto the female cuticle, producing a male “perfume” effect (House *et al*., 2024) that may alter the female’s post-mating chemical phenotype and how she is perceived by subsequent males. In decorated crickets, male-derived CHCs present on females provide information to subsequent males and influence their reproductive behaviour (House *et al*., 2024). Male-derived compounds can also reduce female attractiveness after mating. In butterflies, males transfer anti-aphrodisiac compounds during mating that reduce harassment of recently mated females (Andersson *et al*., 2000). Similarly, in *Drosophila melanogaster*, males transfer anti-aphrodisiac pheromones during copulation, reducing the attractiveness of mated females to subsequent males (Scott, 1986; Laturney & Billeter, 2016).

Because male CHCs can be transferred to females during copulation and alter female signalling, any factor that modifies male chemical profiles or their transfer during mating may also reshape the female post-mating phenotype. Heritable bacterial endosymbionts are strong candidates for such effects, as they are increasingly recognised as important drivers of host chemical phenotypes, mating behaviour and reproductive communication. These endosymbiotic bacteria, including *Wolbachia*, *Cardinium, Rickettsia* and *Spiroplasma*, are widespread in arthropods and best known for affecting host reproduction through cytoplasmic incompatibility (CI), parthenogenesis induction, feminisation and male killing (Werren *et al*., 2008; Engelstädter & Hurst, 2009). Beyond these well-established reproductive effects, endosymbionts are increasingly recognised as influential players in mating behaviour and sexual communication, with documented effects on courtship success (de Crespigny *et al*., 2006; Schneider *et al*., 2019), mate choice (Miller *et al*., 2010; Tourani *et al*., 2026) and sperm competition (de Crespigny & Wedell, 2006). Yet these effects have mostly been interpreted in terms of pre-mating discrimination or reproductive incompatibility (Miller *et al*., 2010; Tourani *et al*., 2024). Less attention has been paid to whether heritable endosymbionts influence post-mating interactions by altering female chemical cues after copulation. This could occur either through male-derived compounds transferred during mating, or through changes in the female’s own CHC profile after mating. Such effects could influence female attractiveness to later males and reveal an additional role for endosymbionts in sexual communication.

Kelly’s citrus thrips, *Pezothrips kellyanus*, provides a particularly powerful system in which to test endosymbiont effects on mating behaviour and chemical communication (Tourani *et al*., 2024; Tourani *et al*., 2026). This haplodiploid pest naturally carries *Cardinium* (C), which is fixed across populations, whereas *Wolbachia* occurs at variable prevalence and only in association with *Cardinium* (CW). In this host, *Wolbachia* induces CI when CW males mate with either C females or experimentally cured endosymbiont-free (U) females. In contrast, *Cardinium*-induced CI is only evident when C males mate with U females (Nguyen *et al*., 2017); however, U populations have not been found in the field (Nguyen et al., 2016; Katlav *et al*., 2024). In addition to CI induction, *Cardinium* has also been associated with host fitness benefits, including increased body size and stress tolerance, whereas *Wolbachia* occurring in the CW association causes moderate costs to the host (Katlav *et al*., 2022a; Katlav *et al*., 2024; Nguyen *et al*., 2017). In response to the CI imposed by CW males, C females have evolved pre-copulatory avoidance of CW males (Tourani *et al*., 2024), a discrimination mediated by the distinctly different CHC profiles of CW males, most notably due to their exclusive production of tridecane (Tourani *et al*., 2026). This host-endosymbionts system therefore already demonstrates that endosymbiont associations can reshape male chemical phenotype and female behavioural response before copulation. However, an unresolved question is whether there are effects that also persist after mating, e.g. by altering the female chemical profile, and influence later male behaviour. Addressing this question is important because it tests whether endosymbiont-associated male chemistry influences not only mate choice before mating, but also female attractiveness after mating.

Here, we tested whether males leave a chemical “perfume” on females after mating and whether first-male endosymbiont association influences subsequent mating interactions. Specifically, we tested whether a male’s endosymbiont-modified CHC profile can alter the female’s CHC blend after copulation and whether male endosymbiont association affects her attractiveness to subsequent males. We predicted that mating with CW males, whose CHC profiles differ from those of C and U males, would produce a distinctive post-mating CHC signature, including the appearance of tridecane, and reduce female attractiveness to subsequent C males. By doing so, we examined whether heritable endosymbionts can contribute to paternity protection through female chemical remodelling, revealing a previously overlooked route by which microbes may influence sexual selection and their own spread.

## Methods

### Study system and insect endosymbiont lines

Laboratory colonies of *P. kellyanus* were established from individuals collected from a citrus orchard in Kulnura, New South Wales, Australia (33.28° S, 151.22° E) in 2017 (Katlav *et al*., 2022a). Colonies were maintained at 20 ± 1°C, 70 ± 5% relative humidity, and a 16:8 h light:dark photoperiod on organic oranges or lemons supplemented with *Typha* sp. pollen every second day (Katlav *et al*., 2021a, 2023; Nguyen *et al*., 2017). Three genetically related lines differing in endosymbiont status were used: a *Cardinium* and *Wolbachia* line (CW) established with field-collected individuals, a *Cardinium*-only line (C), and an endosymbiont-free line (U). The C and U lines were derived from the CW line by antibiotic treatment (tetracycline and rifampicin, respectively) (Katlav *et al*., 2022b). A *Wolbachia*-only line could not be established, consistent with earlier evidence that *Cardinium* provides fitness benefits to the host, while *Wolbachia* imposes fitness costs (Katlav *et al*., 2022a; Nguyen *et al*., 2017). To reduce host genetic variability, CW and C females were introgressed with U males for four generations, followed by multiple generations of maintenance before experiments. Endosymbiont status was confirmed by PCR using *Cardinium*-and *Wolbachia*-specific primers before and after the experiment (Nguyen *et al*., 2016; Tourani *et al*., 2024).

Age-controlled cohorts of virgins were obtained by synchronized oviposition. For each line, approximately 160 females were placed in groups of 20 on individual oranges inside 700 ml plastic containers lined with filter paper. Females oviposited for 48 h and were then removed. Offspring density was kept at approximately 60 larvae per orange to minimize density-dependent effects on development (Katlav *et al*., 2021b). Female and male pupae were separated by their obvious size dimorphism and kept in Petri dishes (35 mm diameter, 10 mm height) with moistened filter paper until adult emergence. All experimental thrips were used as virgins at 1-2 days post-eclosion.

### Experimental design and sequential mating assays

To test whether first-male endosymbiont association altered female chemical profiles after mating and subsequent attractiveness, we conducted sequential mating assays linked with analyses of female CHCs. Virgin females of each endosymbiont association (U, C, and CW) were first mated with a virgin male of one of the three endosymbiont associations (U, C, or CW), generating nine first-mating treatments. All second males used in the behavioural assays were virgin C males because our hypothesis specifically concerned whether mating with a CW male reduces subsequent female attractiveness to C males. Using a single second-male type also standardised male endosymbiont association across treatments.

First matings were conducted individually in mating arenas made from wells of Corning 96-well CellBIND microplates. Each well (6 mm diameter, 10 mm height) contained one virgin female and one virgin male of the designated endosymbiont associations and was covered with cellophane wrap to prevent escape, with a small pinhole made in the wrap for ventilation. Females were monitored under a stereomicroscope until copulation. After copulation, each mated female was removed and held individually for 24 h in a Petri dish (30 mm diameter, 15 mm height) without oviposition substrate, with access to 50% (w/v) honey water and *Typha* sp. pollen.

One day after the first mating, each female was transferred to a clean mating arena for 5 min and then presented with a virgin C male. A previous study used a 5-day interval between the first and second mating because it focused on sperm precedence and found the highest female remating at 5 days after mating (Tourani *et al*., in press). Here, we used a 24 h interval because females may begin laying eggs before 5 days after mating, and our laboratory observations suggest that remating is uncommon once oviposition has started. This allowed us to test whether first-male effects on female attractiveness were already detectable during the early post-mating period. Female-male interactions were observed for 10 min and scored as a sequential process: (1) whether the second male made antennal contact with the female and (2) conditional on contact, whether the male initiated a mating attempt. This design allowed us to distinguish whether the first male affected the chance that a second male detected or contacted the female, or whether it affected the second male’s decision to attempt mating after contact had occurred. Each behavioural treatment combination included 15 replicate females per experimental block. The assay was repeated across three independent experimental blocks, each established from a separate thrips generation using newly synchronized age-matched cohorts. Across the three blocks, this gave a total of 45 replicate females per treatment combination.

### Female CHC extraction and GC-MS analysis

To test whether first-male endosymbiont association altered female CHC profiles, CHCs were extracted from virgin and mated females across treatment groups (Table S1). For each treatment, virgin females were maintained without male exposure, whereas mated females were obtained by placing 50 virgin females and 50 virgin males of the designated endosymbiont type together in a 700 ml plastic container for 24 h to ensure high levels of mating. Both sexes were then freeze-killed at –20 °C, and females were collected for CHC extraction. Virgin females were processed in parallel but without male exposure. For each treatment, ∼100 females were pooled per sample to obtain sufficient material for GC-MS analysis. Each treatment included 22–26 biological replicates across the three experimental blocks. Each block was established from a separate thrips generation using newly synchronised age-matched cohorts.

CHCs were extracted by immersing pooled females in 50 µl of n-hexane. Samples were gently agitated at room temperature for 15 min, after which the solvent was transferred to glass vials with reduced-volume inserts and evaporated under a gentle stream of nitrogen gas. Samples were reconstituted in 30 µl of *n*-hexane containing dodecane at 10 ng/µl, which was used as a post-extraction response standard. Samples were analysed on an Agilent 7890A GC coupled to a 5975C MSD fitted with a DB-5MS capillary column (30 m × 0.25 mm ID, 0.25 µm film thickness). The oven program began at 60 °C for 1 min, increased at 10 °C min⁻¹ to 200 °C, then at 3 °C min⁻¹ to 300 °C, and held for 10 min. Helium was used as the carrier gas at 1.0 ml min⁻¹. Mass spectra were acquired in electron ionization mode at 70 eV over m/z 50–550. Compounds were identified from diagnostic fragmentation patterns and retention indices relative to *n*-alkane standards (C9– C40) (Carlson & Brenner, 1988), and peak areas were integrated in Agilent MassHunter.

## Statistical analyses

Behavioural responses were analysed as a sequential process using binomial logistic regression. The first model tested the probability of antennal contact by the second male, and a second model tested the probability of mating attempt conditional on contact occurring. Because all second males were C, these models tested how female endosymbiont association and first-male endosymbiont association influenced female attractiveness to a C male. Both models included female endosymbiont association (U, C, CW), first-male endosymbiont association (U, C, CW), their interaction and experimental block as fixed effects. Significance of model terms was assessed using likelihood-ratio tests. Because the interaction term was not significant in either model, the main effect of first-male type was interpreted from additive models. Pairwise contrasts among first-male types were reported as odds ratios (OR) with 95% confidence intervals, and pooled response proportions were summarised with Wilson 95% binomial confidence intervals.

CHC peak areas were normalised to the post-extraction dodecane response standard and log10-transformed prior to analysis. Because zeros for tridecane represented true absence rather than valid continuous abundance values, tridecane was excluded from the continuous multivariate CHC matrix and analysed separately. Experimental block was checked initially as a potential source of variation in the female CHC dataset but did not explain significant multivariate structure (Pillai’s trace = 0.169, F = 0.90, p = 0.663). Because no significant experimental block effect was detected, data were pooled for the subsequent treatment-based analyses. The remaining female CHC compounds were then analysed across the 10 female treatments (three virgins and seven mating treatments) using discriminant function analysis (DFA) on column-values. Treatment separation was visualised on the first two discriminant axes. Following House *et al*. (2024), sequential Wilks’ lambda tests were used to assess the significance of the discriminant functions, and classification success was evaluated using leave-one-out cross-validation (LOOCV), whereby each sample was removed once, classified using a DFA model fitted to the remaining samples and compared with its known treatment class. To identify which CHC compounds contributed most to treatment discrimination, we examined the structure matrix and considered compounds with absolute loadings of |0.30| or greater as biologically meaningful contributors.

Differences among treatments in DFA scores were tested by one-way ANOVA for DF1 and DF2 separately, followed by Tukey post-hoc comparisons. Although all treatment classes were included in the analysis, interpretation focused on contrasts relevant to the biological questions of whether mating altered female CHC phenotype relative to virgins and whether male symbiont status altered female CHC phenotype within female background. Geometric distances among treatment centroids were calculated across all discriminant functions to quantify multivariate separation among treatment means.

Tridecane was analysed separately in two ways. First, detection across the 10 female treatments was tested using a chi-square contingency test, and treatment-specific detection proportions were summarised with Wilson 95% binomial confidence intervals. Second, among the three treatments in which tridecane was detected (U_xCW_, C_xCW_ and CW_xCW_ representing U, C and CW females mated with CW males, respectively), abundance was analysed by one-way ANOVA on the log_10_-transformed standardised values, followed by Tukey post-hoc tests. Detected-only abundance is reported as mean ± SE in tables and shown with 95% confidence intervals in figures. All analyses were conducted in R 4.3.0 (R Core Team, 2023).

## Results

### Mating with CW males reduces female attractiveness

First-male endosymbiont association had a strong effect on the first contact by a subsequent C male (LR χ² = 58.55, df = 2, *p* <0.001), whereas female endosymbiont association (LR χ² = 1.54, df = 2, *p* = 0.463), experimental block (LR χ² = 0.03, df = 2, p = 0.984) and the female × first-male endosymbiont interaction (LR χ² = 7.52, df = 4, *p* = 0.111) were not significant (Table 1). Females first mated with a CW male were contacted less frequently by a C male (51.9%) than females first mated with a C male (88.9%) or a U male (85.2%; Table 2). Pairwise contrasts confirmed that a first mating with a CW male significantly reduced the odds of subsequent contact relative to a first mating with a C male (OR = 0.13, 95% CI = 0.07–0.25, *p* <0.001) or a U male (OR = 0.19, 95% CI = 0.10–0.33, *p* <0.001), whereas the first mating with C and U males did not differ when the second mating was with a C male (OR = 0.72, 95% CI = 0.35–1.47, *p* = 0.366; Table 3).

**Table 1.**
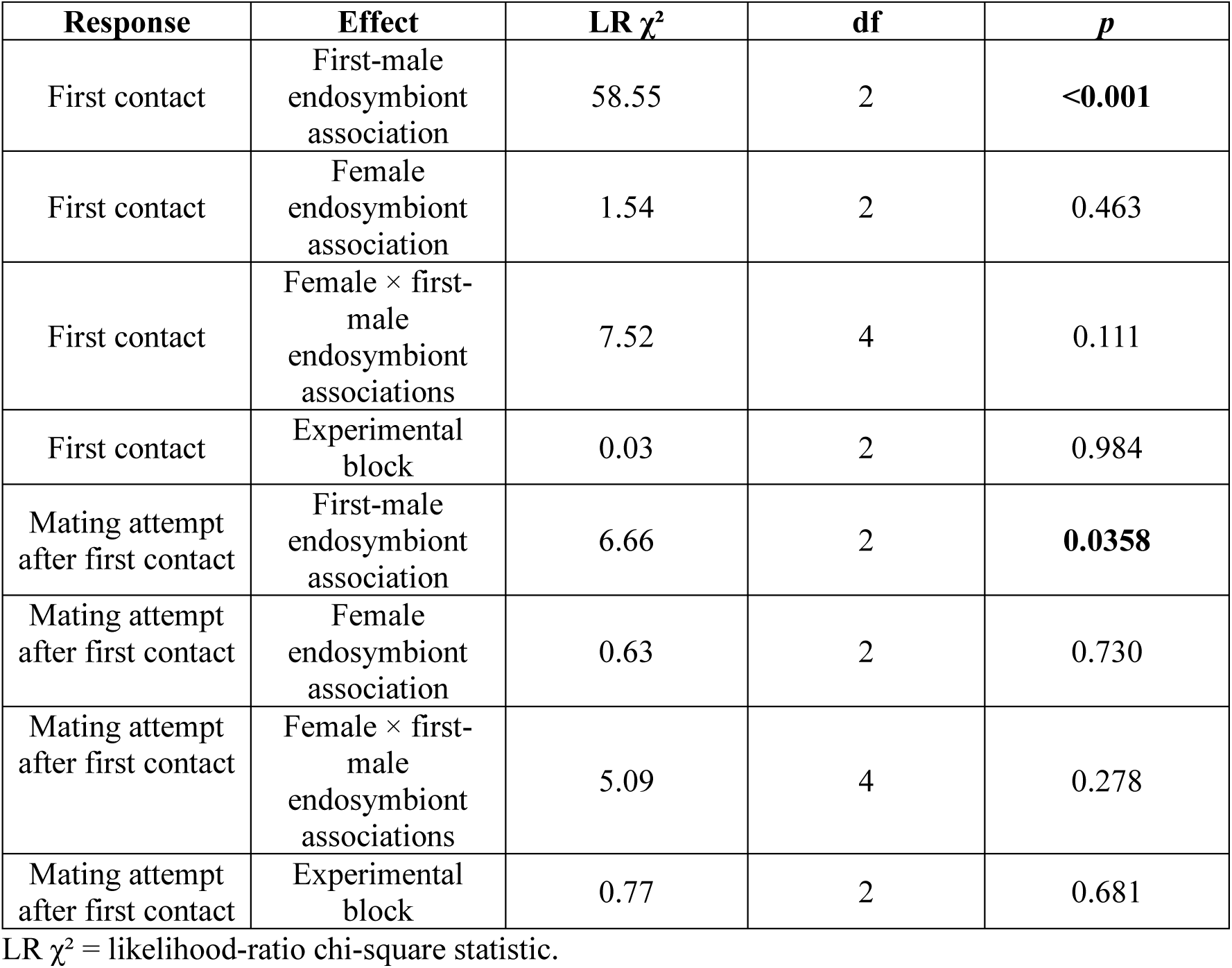
Binomial logistic model effects of female endosymbiont association, first-male endosymbiont association and experimental block on female attractiveness to a *Cardinium*-carrying second male.

| Response | Effect | LR $\chi^2$ | df | *p* |
| --- | --- | --- | --- | --- |
| First contact | First-male endosymbiont association | 58.55 | 2 | **<0.001** |
| First contact | Female endosymbiont association | 1.54 | 2 | 0.463 |
| First contact | Female × first-male endosymbiont associations | 7.52 | 4 | 0.111 |
| First contact | Experimental block | 0.03 | 2 | 0.984 |
| Mating attempt after first contact | First-male endosymbiont association | 6.66 | 2 | **0.0358** |
| Mating attempt after first contact | Female endosymbiont association | 0.63 | 2 | 0.730 |
| Mating attempt after first contact | Female × first-male endosymbiont associations | 5.09 | 4 | 0.278 |
| Mating attempt after first contact | Experimental block | 0.77 | 2 | 0.681 |

LR $\chi^2$ = likelihood-ratio chi-square statistic.

**Table 2.** Pooled probabilities of first contact and mating attempt by a *Cardinium*-carrying second male after female first mating with endosymbiont-free (U), *Cardinium*-only (C), or *Cardinium*-*Wolbachia* (CW) male.

| First-male endosymbiont | First contact (successful males/total males) | Proportion % (95% CI) | Mating attempt after first contact (successful males/total males) | Proportion % (95% CI) |
| --- | --- | --- | --- | --- |
| U | 115/135 | 85.2 (78.2–90.2) | 69/115 | 60.0 (50.9–68.5) |
| C | 120/135 | 88.9 (82.5–93.1) | 73/120 | 60.8 (51.9–69.1) |
| CW | 70/135 | 51.9 (43.5–60.1) | 30/70 | 42.9 (31.9–54.5) |

**Table 3.** Pairwise odds-ratio contrasts among first-male endosymbiont types from additive binomial logistic models of first contact and mating attempt by a *Cardinium*-carrying second male.

| Response | Contrast | Odds ratio (95% confidence interval) | *p* |
| --- | --- | --- | --- |
| First contact | U vs C | 0.72 (0.35–1.47) | 0.366 |
| First contact | CW vs C | 0.13 (0.07–0.25) | **<0.001** |
| First contact | CW vs U | 0.19 (0.10–0.33) | **<0.001** |
| Mating attempt after first contact | U vs C | 0.96 (0.57–1.63) | 0.887 |
| Mating attempt after first contact | CW vs C | 0.48 (0.26–0.88) | **0.0176** |
| Mating attempt after first contact | CW vs U | 0.50 (0.27–0.92) | **0.0253** |

A similar but weaker pattern was observed for mating attempts by a subsequent male after a first contact. Again, first male endosymbiont association was significant (LR χ² = 6.66, df = 2, *p* = 0.0358), whereas female endosymbiont association (LR χ² = 0.63, df = 2, *p* = 0.730), experimental block (LR χ² = 0.77, df = 2, *p* = 0.681) and the interaction (LR χ² = 5.09, df = 4, *p* = 0.278) were not significant (Table 1). Again, females first mated with a CW male were less attractive to a second C male, eliciting mating attempts in only 42.9% of contacted trials, compared with 60.8% after first mating with a C male and 60.0% after first mating with a U male (Table 2). Pairwise contrasts showed that first mating with a CW male reduced the odds of a subsequent mating attempt by a C male relative to first mating with a C male (OR = 0.48, 95% CI = 0.26–0.88, *p* = 0.0176) or a U male (OR = 0.50, 95% CI = 0.27–0.92, *p* = 0.0253), whereas the C and U first mating treatments again did not differ (OR = 0.96, 95% CI = 0.57–1.63, *p* = 0.887; Table 3).

### Female CHC profiles are changed by mating, with CW males producing a distinctive post-mating signature

The tridecane-excluded female CHC profiles were clearly different among all female virgin and mating treatments (Wilks’ lambda = 8.89 × 10⁻⁵, χ² = 2126.86, df = 198, *p* < 0.001) (Fig. 1). In the tridecane-excluded DFA matrix of CHC profiles, DF1 and DF2 explained 60.0% and 21.4% of the among-treatment separation, respectively. Cross-validation (LOOCV) correctly assigned 84.9% of samples to their treatment class (Fig. 1, Table S2). Classification accuracy was highest for C females, reaching 100% for C_V_, C_xC_, and C_xCW_, and 95.7% for C_xU_, whereas discrimination was weaker among some U female treatments, particularly U_xU_ and U_xC_ (Table S2). Mating treatment effects were highly significant on both DF1 (F_9,235_= 602.592, *p* < 0.001) and DF2 (F_9,235_= 215.325, *p* < 0.001) (Table S3). To interpret the DFA axes, we examined structure coefficients for individual CHCs on DF1 and DF2 (Table 4). Except for heptadecane, DF1 was positively associated with all CHCs, indicating that the primary separation among treatments was driven largely by variation in overall CHC abundance, with higher DF1 scores corresponding to greater CHC abundance (Table 4). This dimension largely separates C females from U and CW females (Fig. 1). DF2 was positively associated with six compounds (heptadecane, 7-methylheptadecane, tricosane, heptacosane, nonacosane and hentriacontane) and negatively associated with 11-methyltricosane and pentacosane (Table 4). DF2 primarily separated treatments according to first-male endosymbiont association: females mated with CW males generally had higher scores, whereas females mated with U or C males had lower scores, with virgin females occupying intermediate positions (Fig. 1).

**Fig. 1.**
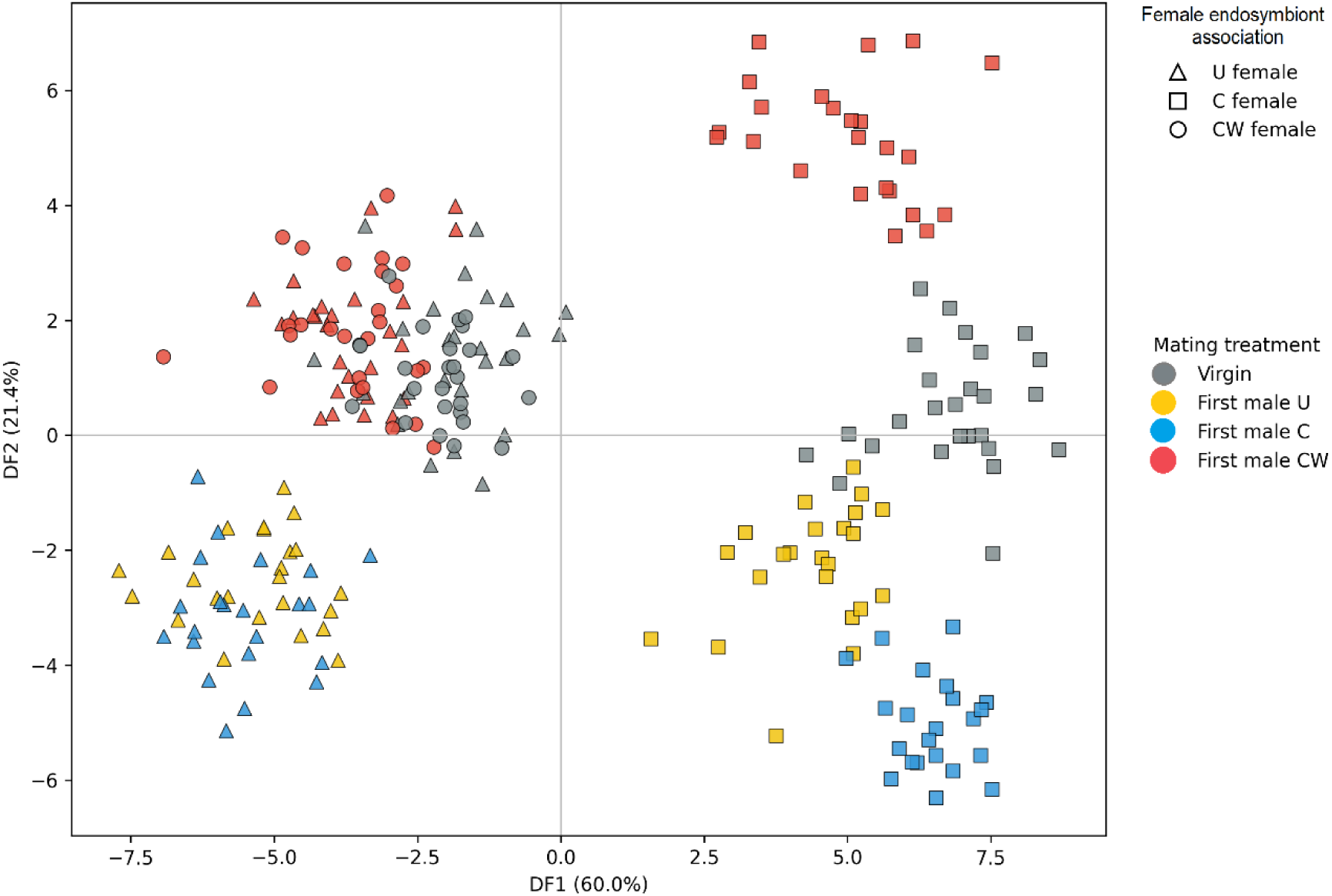
Discriminant function analysis of female cuticular hydrocarbon profiles across all tested mating treatments of females with different endosymbiont associations (tridecane excluded). The CHC experiment used a reduced treatment design informed by the behavioural results. Behavioural assays did not indicate that mating with U or C males altered the subsequent attractiveness of CW females to a second C male. We therefore did not include CW_xU_ and CW_xC_ treatments in the CHC analysis. However, U and C males were retained in crosses with U and C females as control treatments, allowing us to compare these baseline mating effects with the effect of mating with CW males.

**Table 4.** Structure coefficients for discriminant function analysis (DFA) of female cuticular hydrocarbon (CHC) profiles across all female treatments.

| CHC compounds | DF1 | DF2 |
| --- | --- | --- |
| Heptadecane | 0.074 | **0.765** |
| 7-methyl-heptadecane | **0.320** | **0.478** |
| Nonadecane | **0.392** | -0.176 |
| Heneicosane | **0.449** | 0.169 |
| Tricosane | **0.631** | **0.394** |
| 11-methyltricosane | **0.586** | **-0.405** |
| Tetracosane | **0.617** | 0.283 |
| 2-methyltetracosane | **0.782** | 0.290 |
| Pentacosane | **0.797** | **-0.300** |
| 11-methylpentacosane | **0.757** | 0.199 |
| 1-hexacosene | **0.732** | 0.253 |
| Heptacosane | **0.801** | **0.347** |
| 13-methylheptacosane | **0.883** | 0.123 |
| Octacosane | **0.930** | 0.180 |
| 2-methyl-octacosane | **0.968** | 0.073 |
| Nonacos-1-ene | **0.753** | 0.215 |
| Nonacosane | **0.761** | **0.404** |
| 15-methylnonacosane | **0.910** | 0.116 |
| Hentriacontane | **0.654** | **0.381** |
| 3-methyldotriacontane | **0.801** | 0.221 |
| Tritriacontane | **0.897** | 0.186 |
| 2-methyl-tritriacontane | **0.453** | 0.278 |

The DFA was conducted without tridecane, which was absent from most female treatments. Structure coefficients with absolute values of 0.30 or greater were considered biologically meaningful contributors and are shown in bold.

Relative to virgins, mating altered female CHC phenotypes for each female endosymbiont association type, although the strength of this effect varied among different mating treatments. In endosymbiont-free females, U females mated with U males (U_xU_) and C males (U_xC_) differed from virgin U females (U_V_) on both DF1 and DF2, whereas U_xCW_ differed from U_V_ on DF1 but not DF2 (Fig. S1, Fig. S2; Table S4, Table S5). In *Cardinium* females, C_xU_ and C_xCW_ differed from C_V_ on both DF axes, whereas C_xC_ differed from C_V_ on DF2 but not DF1 (Fig. S1, Fig. S2, Table S4, Table S5). For CW females, CW_xCW_ differed from CW_V_ on DF1 but not DF2 (Fig. S1, Fig. S2, Table S4, Table S5). Centroid distances also showed clear separation between virgin and mated classes within each female background (Table 5).

**Table 5.** Full matrix of geometric distances among female-treatment centroids across all discriminant functions, based on the tridecane-excluded female cuticular hydrocarbon dataset.

| Treatment | U<sub>V</sub> | U<sub>xU</sub> | U<sub>xC</sub> | U<sub>xCW</sub> | C<sub>V</sub> | C<sub>xU</sub> | C<sub>xC</sub> | C<sub>xCW</sub> | CW<sub>V</sub> | CW<sub>xCW</sub> |
| --- | --- | --- | --- | --- | --- | --- | --- | --- | --- | --- |
| U<sub>V</sub> | - |  |  |  |  |  |  |  |  |  |
| U<sub>xU</sub> | 6.615 | - |  |  |  |  |  |  |  |  |
| U<sub>xC</sub> | 6.824 | 1.744 | - |  |  |  |  |  |  |  |
| U<sub>xCW</sub> | 3.566 | 5.527 | 6.185 | - |  |  |  |  |  |  |
| C<sub>V</sub> | 9.922 | 12.895 | 13.140 | 11.515 | - |  |  |  |  |  |
| C<sub>xU</sub> | 8.052 | 10.001 | 10.110 | 9.626 | 4.788 | - |  |  |  |  |
| C<sub>xC</sub> | 11.417 | 13.250 | 13.367 | 12.657 | 9.317 | 6.947 | - |  |  |  |
| C<sub>xCW</sub> | 8.789 | 13.123 | 13.655 | 9.712 | 7.148 | 8.270 | 10.940 | - |  |  |
| CW<sub>V</sub> | 3.768 | 5.710 | 6.287 | 3.636 | 9.679 | 7.782 | 11.823 | 9.075 | - |  |
| CW<sub>xCW</sub> | 4.295 | 6.058 | 6.518 | 2.808 | 11.932 | 9.901 | 12.605 | 9.697 | 4.903 | - |

First-male endosymbiont association strongly shaped post-mating female CHC composition, with females mated with CW males showing the most distinctive profiles within both U and C female backgrounds. Among endosymbiont-free U females, U_xU_ and U_xC_ were not separated on either DF1 or DF2 (Fig. S1, Fig. S2, Table S4, Table S5), and their centroid distance was small (1.744; Table 5). By contrast, U_xCW_ differed from both U_xU_ and U_xC_ on both axes (Fig. S1, Fig. S2, Table S4, Table S5), with substantially larger centroid distances (5.527 and 6.185, respectively; Table 5). A similar pattern was evident in C females. Although C_xU_ and C_xC_ were separated from one another, C_xCW_ was strongly differentiated from both C_xU_ and C_xC_, with centroid distances of 8.270 and 10.940, respectively, compared with 6.947 between C_xU_ and C_xC_ (Table 5). Consistent with this, C_xCW_ differed from C_xC_ on both DF1 and DF2, and from C_xU_ on DF2 (Fig. S1, Fig. S2, Table S4, Table S5).

Tridecane provided the clearest single-compound signature of this effect. It was absent from all virgin females and from females mated with U or C males, but was present in females mated with CW males, producing a strong treatment effect on detection (χ² = 245.0, df = 9, *p* < 0.001; Table S6, Fig. 2A). Among mated females from which tridecane was detected, its abundance also differed significantly (F = 1493.206, *p* < 0.001; Table 6, Fig. 2B), with the highest mean in C_xCW_ females (1.647 ± 0.009 SE), intermediate mean in U_xCW_ females (1.129 ± 0.012 SE), and the lowest mean in CW_xCW_ females (0.887 ± 0.008 SE). Together, these results identify a CW-associated post-mating signature comprising a distinct multivariate CHC phenotype and the exclusive occurrence of tridecane.

**Fig. 2.**
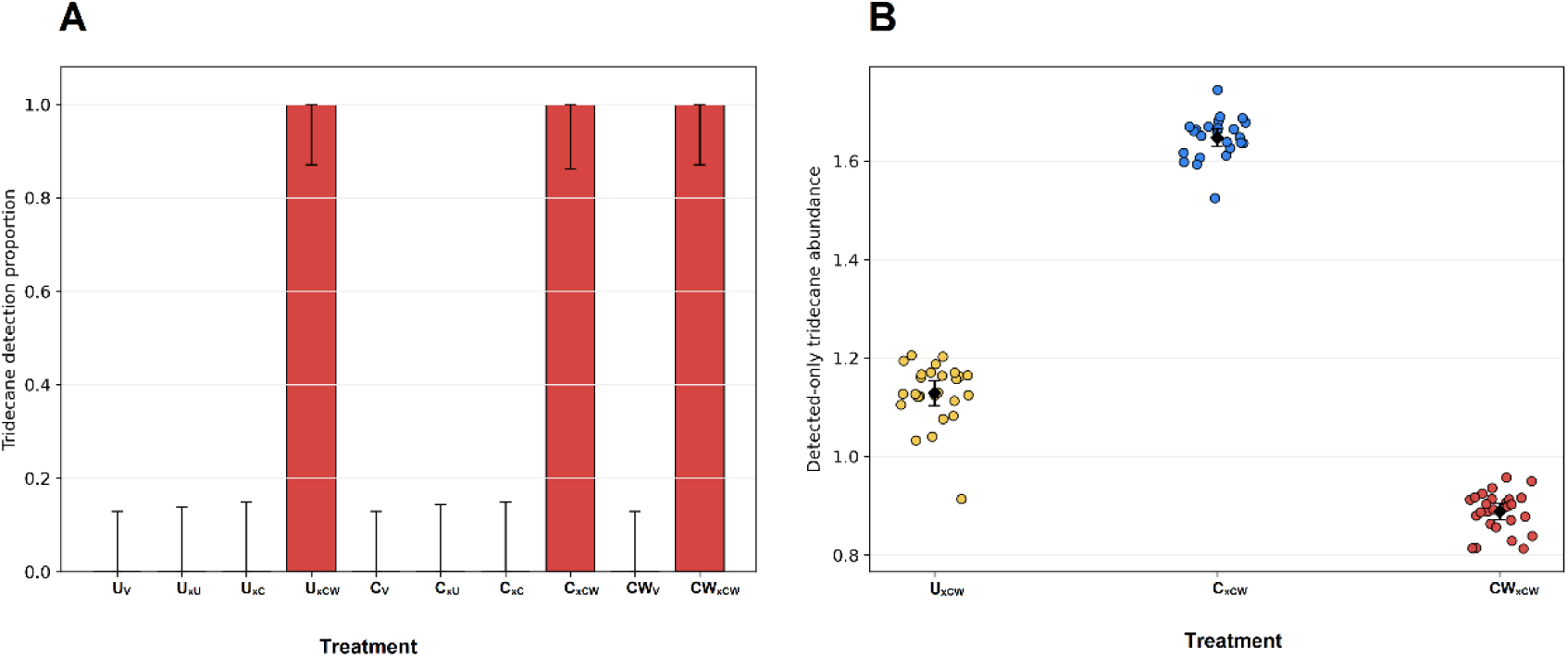
Occurrence and abundance of tridecane in female cuticular hydrocarbon profiles. **(A)** Treatment-specific proportion of females with detectable tridecane. Bars show the proportion of pooled-female samples in which tridecane was detected, and error bars show Wilson 95% binomial confidence intervals for detection probability. The confidence intervals are shown because tridecane detection is a binary response, and they indicate uncertainty around the estimated detection proportion. **(B)** Detected-only tridecane abundance in U_xCW_, C_xCW_, and CW_xCW_ females. Points represent independent pooled-female samples, black symbols show group means, and error bars indicate 95% confidence intervals.

**Table 6.**
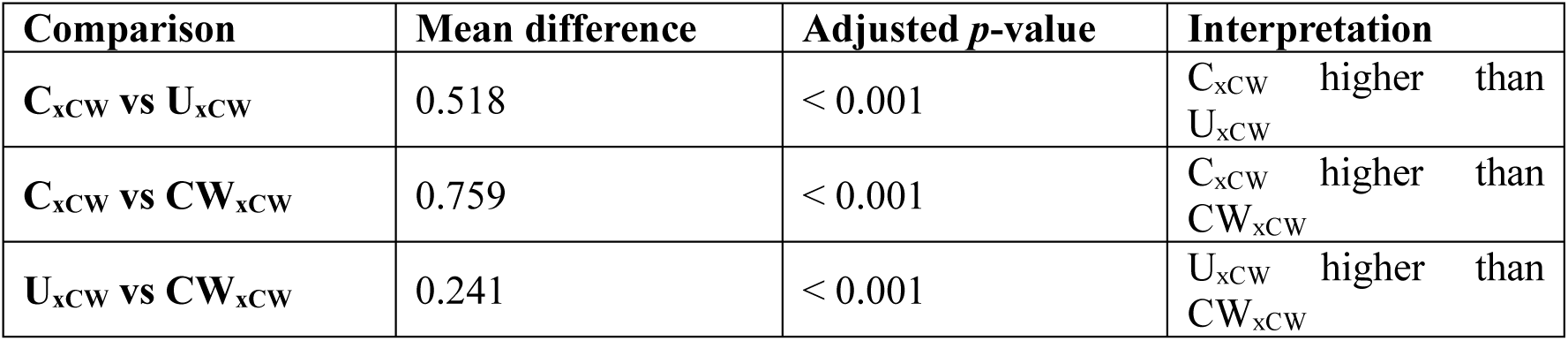
Pairwise Tukey comparisons of tridecane abundance among females mated with CW males (U_xCW_, C_xCW_ and CW_xCW_).

| Comparison | Mean difference | Adjusted *p*-value | Interpretation |
| --- | --- | --- | --- |
| C<sub>xCW</sub> vs U<sub>xCW</sub> | 0.518 | < 0.001 | C<sub>xCW</sub> higher than U<sub>xCW</sub> |
| C<sub>xCW</sub> vs CW<sub>xCW</sub> | 0.759 | < 0.001 | C<sub>xCW</sub> higher than CW<sub>xCW</sub> |
| U<sub>xCW</sub> vs CW<sub>xCW</sub> | 0.241 | < 0.001 | U<sub>xCW</sub> higher than CW<sub>xCW</sub> |

## Discussion

Our study shows that mating reshapes CHC profiles of *P. kellyanus* females, and that the extent of this change depends strongly on the endosymbiont association of the first male. Females from all mating combinations showed some degree of chemical change relative to virgin females, indicating that mating itself can alter the female CHC phenotype. However, mating with CW males carrying CI-inducing *Wolbachia* produced a distinctive male-associated post-mating signature, most clearly marked by the appearance of tridecane. Females first mated with CW males were less attractive to subsequent C males, receiving fewer antennal contacts and fewer mating attempts after contact. Most notably, tridecane, which was absent from all virgin females and from females mated with U or C males, was detected after females had mated with CW males. This indicates that a chemical compound previously characterised as distinctive of CW males and linked to pre-copulatory discrimination by C females against incompatible CW males (Tourani *et al*., 2026) can also be detected in the female CHC profile after mating. Together, these findings extend earlier work on endosymbiont-mediated mate choice in *P. kellyanus* by showing that endosymbiont-associated chemical effects are not limited to mate recognition before copulation but can also influence the signals females carry after mating.

### Symbiont-associated chemistry extends sexual communication beyond mating

Our study shows that mating changes a female’s chemical profile irrespective of her endosymbiont association. However, the clearest male-specific qualitative change occurred after mating with CW males: tridecane, which was absent from virgin females and females mated with U or C males, was detected after mating with CW males. Tridecane was previously found as a distinctive compound in CW males and was absent from C and U males (Tourani *et al*., 2026). This indicates that a chemical signature associated with CW males before copulation also appears in the female CHC profile after mating. Importantly, we did not detect any additional CHC compound that appeared only on mated females while being absent from both virgin females and males. This suggests that the clearest qualitative change following mating with a CW male is the acquisition of the CW male-associated compound tridecane, rather than the production of a completely new female-specific CHC after mating. Therefore, endosymbiont-associated male CHC signalling is not only involved in pre-copulatory mate recognition but may also shape the chemical cues displayed by females after mating.

This interpretation fits with the broader role of CHCs as surface compounds involved in waterproofing and chemical communication in insects (Howard & Blomquist, 1982; Blomquist & Bagnères, 2010). It is also consistent with evidence that mating can alter female chemical profiles through contact or transfer of male-derived compounds, as reported in crickets (House *et al*., 2024) and *Drosophila* (Laturney & Billeter, 2016). In *Drosophila*, cis-vaccenyl acetate (cVA) provides a well-studied example: it is a male-derived pheromone transferred to females during copulation, contributes to reduced courtship toward mated females (Ejima *et al*., 2007) and is detected through Or67d-expressing olfactory neurons that mediate behavioural responses to cVA (Kurtovic *et al*., 2007).

The multivariate structure of female CHC profiles suggests that symbiont background and mating history shape different dimensions of the chemical phenotype. DF1 primarily captured quantitative variation in overall CHC abundance, with *Cardinium*-associated females occupying a high-abundance region distinct from endosymbiont-free and *Cardinium–Wolbachia*-associated females. Because this divergence was already present among virgins, it most likely reflects female physiological background rather than male-derived transfer. Endosymbiont-associated changes in host metabolism, resource allocation or hydrocarbon biosynthesis could underlie this pattern, although its functional significance remains unresolved. DF2, in contrast, reflected compositional restructuring of the blend. The pronounced separation of C_xCW_ females from C females mated with U or C males indicates that the post-mating phenotype depends on the interaction between female and male symbiont backgrounds rather than representing a uniform mating response. This context dependence is difficult to reconcile with simple passive coating alone and instead suggests selective retention, differential grooming or female-mediated remodelling of the blend. The C_xCW_ phenotype may therefore represent an interaction-dependent chemical state through which an incompatible mating alters the female’s subsequent sexual phenotype, providing a plausible mechanism linking symbiont-associated reproductive incompatibility to later male discrimination.

Tridecane is especially informative in this context because it provides a chemically discrete marker embedded within the broader CW-associated shift. Across insects, tridecane occurs in both defensive and communicative systems: it contributes to defensive secretions and escape responses in heteropterans and beetles (Eisner *et al*., 1977; Krall *et al*., 1999; Zarbin *et al*., 2000), acts as a pheromonal component in some Hymenoptera (Xu *et al*., 2023), and elicits defensive behaviour in the thrips *Liothrips jatrophae* (González-Orellana *et al*., 2022). In *P. kellyanus*, the restriction of tridecane to CW males and to females after mating with CW males makes it a plausible cue of previous-male identity or mating history. Its highest abundance in C_xCW_ females, which occupied a distinct region of multivariate CHC space relative to C_xU_ and C_xC_ females, suggests that tridecane forms part of a broader post-mating signature. Together with its association with sexual discrimination in this species (Tourani *et al*., 2026), this pattern supports a role in reducing subsequent male engagement. However, causality remains unresolved because males may respond to tridecane itself, its relative abundance, or the entire blend. Manipulative assays using reconstructed CHC blends are needed to determine whether males respond specifically to tridecane or to the broader post-mating chemical profile.

### Post-mating chemical remodelling reduces female attractiveness

The results of the behavioural experiments indicate that the change in female CHC profile after mating with a CW male affected how a second approaching male responded to the female. Females first mated with CW males received fewer antennal contacts from subsequent C males, and such females were less likely to receive mating attempts. Functionally, this resembles post-mating paternity-protection strategies in which males that mated with a female reduce access to this female by subsequent males or reduce their reproductive success (Parker, 1970; Birkhead & Pizzari, 2002). Such strategies occur across diverse animals; for example, in birds, males may guard females or increase copulation frequency during the fertile period to reduce extra-pair fertilisation (Møller & Birkhead, 1991), while in mammals and other taxa, copulatory plugs can interfere with subsequent sperm transfer or fertilisation success (Dixson & Anderson, 2002; Mangels *et al*., 2016; Sutter & Lindholm, 2016). In insects, males can also reduce female remating or attractiveness through seminal proteins, mating plugs and transferred anti-aphrodisiac compounds (Wigby & Chapman, 2005; Avila *et al*., 2011). Paternity protection via chemical cues is especially relevant here because male-derived compounds can alter how subsequent males may respond to mated females, as shown in *Drosophila* (Laturney & Billeter, 2016) and pierid butterflies (Andersson *et al*., 2000; Andersson *et al*., 2003). Our results suggest a similar post-mating chemical effect in *P. kellyanus*, but with an added microbial component: the chemical signature associated with CW males appears on females after mating and is linked with reduced attractiveness to subsequent males, with potential effects on *Wolbachia* spread in populations.

This effect may be especially important in *P. kellyanus*, where strong first-male sperm precedence means that fertilisation outcomes are shaped primarily by the first mating, although subsequent mating can still alter offspring outcomes (Tourani *et al*., in press). When C females first mated with incompatible CW males, a subsequent compatible mating with C males restored a modest level of female offspring production, although offspring sex ratio remained strongly influenced by the first male (Tourani *et al*., in press). Thus, the previous mating-order study provides strong evidence for the effect of mating order on reproductive outcome, because it measured CI expression, offspring sex ratio and female fitness. Our present study makes a different but complementary contribution by identifying a possible behavioural and chemical mechanism that could influence whether such compatible remating occurs. In this context, a CW-associated post-mating chemical signal that reduces female attractiveness could be reproductively important. If females mated with CW males are less likely to attract or accept later C males, the opportunity for a subsequent mating with a compatible C male may be reduced, potentially preserving the reproductive influence of the first CW male. Together, these findings suggest that post-mating chemical remodelling could contribute to paternity protection in *P. kellyanus*.

### Consequences for *Wolbachia* dynamics in host populations

The post-mating changes in CHCs observed here may help explain why *Wolbachia* dynamics in *P. kellyanus* are more complex than expected from CI alone. Field surveys show that *Cardinium* is fixed across sampled populations, being detected in all populations at near-100% prevalence, whereas *Wolbachia* varies geographically, with high prevalence in coastal populations of New South Wales and Queensland but low prevalence in inland populations of eastern Australia, as well as in Victoria, South Australia and Western Australia, as well as absence in populations of the invasive range in New Zealand and the Mediterranean region (Katlav *et al*., 2024). In addition, *Wolbachia* is associated with reduced mitochondrial diversity in host populations, suggesting a recent or ongoing *Wolbachia*-driven mitochondrial selective sweep in *P. kellyanus* in eastern Australia (Katlav *et al*., 2024). However, laboratory studies suggest that *Wolbachia* spread is shaped by factors beyond maternal transmission and CI strength: *Cardinium* is associated with several host fitness benefits, while *Wolbachia* in the CW background has been linked to fitness costs (Nguyen *et al*., 2017; Katlav *et al*., 2022b; Katlav *et al*., 2024). These patterns indicate that while *Wolbachia* has the potential to spread, its spread may be constrained by costs to the host and by host behaviours that reduce the incidence of incompatible matings (Tourani *et al*., 2024, 2026).

Our study species *P. kellyanus* therefore displays evolutionary conflicts between maternally transmitted endosymbionts and host reproductive interests. Reproductive parasites such as *Wolbachia* can gain a transmission advantage through CI, but CI also creates selection for host counter-adaptations that reduce the costs of incompatible matings, including mate discrimination (Champion de Crespigny *et al*., 2005), reduced receptivity and post-copulatory mechanisms that limit incompatible sperm use (Werren *et al*., 2008; Birkhead & Pizzari, 2002; Rowe *et al*., 2020). In *P. kellyanus*, this conflict may operate at both pre-and post-mating stages. Before mating, the distinct chemical profile of CW males exposes them to avoidance by C females (Tourani *et al*., 2024, 2026). After mating, however, the same male-associated chemistry may reduce female attractiveness to subsequent C males, helping preserve the reproductive effect of the first CW male. Thus, *Wolbachia* dynamics in *P. kellyanus* may reflect an arms race between symbiont-driven reproductive manipulation and host behaviours that limit the costs of CI. Future studies tracking endosymbiont frequencies, mating behaviour and CHC variation in field populations will be needed to test how these processes interact under natural conditions.

## Conclusion

Our findings show that endosymbiont effects on sexual communication in *P. kellyanus* extend beyond mate choice before copulation. Previous studies showed that C females have reduced receptivity to incompatible CW males and that this discrimination is associated with the distinct CHC profile of CW males, which includes tridecane (Tourani *et al*., 2024; Tourani *et al*., 2026). Here, we show that mating changes the females’ CHC profile after mating, and in particular, the CHC profile of C females after mating with incompatible CW males. This the reduces these females’ attractiveness for subsequent compatible C males, a situation that may be encountered in field populations that consist of C and CW individuals. This suggests that symbiont-associated male chemistry can persist beyond copulation and alter the signal of mated females encountered by subsequent males.

In the context of the strong first-male sperm precedence observed in *P. kellyanus*, these post-mating chemical effects may be especially relevant, because they could further reduce the already limited opportunity for a later compatible male to modify reproductive outcomes after an incompatible CW first mating (Tourani *et al*., in press). More broadly, our study helps connect endosymbiont-mediated CI, CHC signalling and remating behaviour in a more holistic framework, where CI and first-male sperm precedence are likely the strongest determinants of reproductive outcome, and post-mating CHC remodelling may further reduce male access to already-mated females. Endosymbionts may therefore shape host reproduction not only through embryonic mortality or pre-copulatory mate choice, but also through post-mating changes in female chemical phenotype. By doing so, they may contribute to paternity protection through female chemical remodelling, revealing a further route by which microbes can influence sexual selection and their own spread. Future work should test whether tridecane itself, or the broader CW-associated CHC blend, directly affects male behaviour, sperm use, paternity outcomes and endosymbiont dynamics under natural conditions.

## Disclosure

The authors declare no competing interests.

