## Supplementary tables & Figures for "Symbiont-mediated shifts in cuticular hydrocarbon profiles reduce female attractiveness after mating"

**Table S1.** Overview of female mating treatments to assess effect of male CHC profile on female CHC profile.

| **Treatment code** | **Female endosymbiont association** | **Female status** | **First male endosymbiont association** | **Meaning** |
| --- | --- | --- | --- | --- |
| U_V_ | Endosymbiont-free | Virgin | - | Endosymbiont-free female, unmated |
| U_xU_ | Endosymbiont-free | Mated | U | Endosymbiont-free female mated with an endosymbiont-free male |
| U_xC_ | Endosymbiont-free | Mated | C | Endosymbiont-free female mated with a *Cardinium*-carrying male |
| U_xCW_ | Endosymbiont-free | Mated | CW | Endosymbiont-free female mated with a *Cardinium*-*Wolbachia* male |
| C_V_ | *Cardinium* | Virgin | - | *Cardinium*-carrying female, unmated |
| C_xU_ | *Cardinium* | Mated | U | *Cardinium*-carrying female mated with an endosymbiont-free male |
| C_xC_ | *Cardinium* | Mated | C | *Cardinium*-carrying female mated with a *Cardinium*- carrying male |
| C_xCW_ | *Cardinium* | Mated | CW | *Cardinium*-carrying female mated with a *Cardinium*-*Wolbachia* male |
| CW_V_ | *Cardinium*-*Wolbachia* | Virgin | - | *Cardinium*-*Wolbachia* female, unmated |
| CW_xCW_ | *Cardinium*-*Wolbachia* | Mated | CW | *Cardinium*-*Wolbachia* female mated with a *Cardinium*-*Wolbachia* male |

**The CHC experiment used a reduced treatment design informed by the behavioural results. Behavioural assays did not indicate that mating with U or C males altered the subsequent attractiveness of CW females to a second C male. We therefore did not include CW_xU_ and CW_xC_ treatments in the CHC analysis. However, U and C males were retained in crosses with U and C females as control treatments, allowing us to compare these baseline mating effects with the effect of mating with CW males.**

**Table S2.** Leave-one-out cross-validated (LOOCV) classification accuracy from discriminant function analysis (DFA) of female cuticular hydrocarbon profiles across all female treatments, excluding tridecane.

| **Treatment** | **n** | **Correctly classified** | **LOOCV accuracy (%)** |
| --- | --- | --- | --- |
| U_V_ | 26 | 22 | 84.6 |
| U_xU_ | 24 | 15 | 62.5 |
| U_xC_ | 22 | 16 | 72.7 |
| U_xCW_ | 26 | 21 | 80.8 |
| C_V_ | 26 | 26 | 100.0 |
| C_xU_ | 23 | 22 | 95.7 |
| C_xC_ | 22 | 22 | 100.0 |
| C_xCW_ | 24 | 24 | 100.0 |
| CW_V_ | 26 | 20 | 76.9 |
| CW_xCW_ | 26 | 20 | 76.9 |

Overall LOOCV classification accuracy = 84.9%. DF1 explained 60.0% and DF2 explained 21.4% of the among-treatment separation.

**Table S3.** One-way ANOVA of discriminant function (DF) scores across female treatments for DF1 and DF2.

| **Discriminant axis** | **Numerator df** | **Denominator df** | **F** | **p** |
| --- | --- | --- | --- | --- |
| DF1 | 9 | 235 | 602.592 | < 0.001 |
| DF2 | 9 | 235 | 215.325 | < 0.001 |

**Table S4.** Tukey post-hoc comparisons of DFA axis 1 scores across all ten female treatments.

| **Comparison** | **Mean difference** | **Adjusted *p*** | **Significant** |
| --- | --- | --- | --- |
| CW_V_ vs CW_xCW_ | 1.5610 | < 0.001 | Yes |
| CW_V_ vs C_V_ | -8.9091 | < 0.001 | Yes |
| CW_V_ vs C_xC_ | -8.5690 | < 0.001 | Yes |
| CW_V_ vs C_xCW_ | -7.1038 | < 0.001 | Yes |
| CW_V_ vs C_xU_ | -6.4410 | < 0.001 | Yes |
| CW_V_ vs U_V_ | -0.2280 | 0.9982 | No |
| CW_V_ vs U_xC_ | 3.4152 | < 0.001 | Yes |
| CW_V_ vs U_xCW_ | 1.5714 | < 0.001 | Yes |
| CW_V_ vs U_xU_ | 3.2582 | < 0.001 | Yes |
| CW_xCW_ vs C_V_ | -10.4701 | < 0.001 | Yes |
| CW_xCW_ vs C_xC_ | -10.1300 | < 0.001 | Yes |
| CW_xCW_ vs C_xCW_ | -8.6648 | < 0.001 | Yes |
| CW_xCW_ vs C_xU_ | -8.0020 | < 0.001 | Yes |
| CW_xCW_ vs U_V_ | -1.7890 | < 0.001 | Yes |
| CW_xCW_ vs U_xC_ | 1.8542 | < 0.001 | Yes |
| CW_xCW_ vs U_xCW_ | 0.0104 | 1.0000 | No |
| CW_xCW_ vs U_xU_ | 1.6972 | < 0.001 | Yes |
| C_V_ vs C_xC_ | 0.3401 | 0.9756 | No |
| C_V_ vs C_xCW_ | 1.8054 | < 0.001 | Yes |
| C_V_ vs C_xU_ | 2.4681 | < 0.001 | Yes |
| C_V_ vs U_V_ | 8.6812 | < 0.001 | Yes |
| C_V_ vs U_xC_ | 12.3244 | < 0.001 | Yes |
| C_V_ vs U_xCW_ | 10.4805 | < 0.001 | Yes |
| C_V_ vs U_xU_ | 12.1674 | < 0.001 | Yes |
| C_xC_ vs C_xCW_ | 1.4652 | < 0.001 | Yes |
| C_xC_ vs C_xU_ | 2.1279 | < 0.001 | Yes |
| C_xC_ vs U_V_ | 8.3410 | < 0.001 | Yes |
| C_xC_ vs U_xC_ | 11.9842 | < 0.001 | Yes |
| C_xC_ vs U_xCW_ | 10.1404 | < 0.001 | Yes |
| C_xC_ vs U_xU_ | 11.8272 | < 0.001 | Yes |
| C_xCW_ vs C_xU_ | 0.6627 | 0.4118 | No |
| C_xCW_ vs U_V_ | 6.8758 | < 0.001 | Yes |
| C_xCW_ vs U_xC_ | 10.5190 | < 0.001 | Yes |
| C_xCW_ vs U_xCW_ | 8.6752 | < 0.001 | Yes |
| C_xCW_ vs U_xU_ | 10.3620 | < 0.001 | Yes |
| C_xU_ vs U_V_ | 6.2131 | < 0.001 | Yes |
| C_xU_ vs U_xC_ | 9.8563 | < 0.001 | Yes |
| C_xU_ vs U_xCW_ | 8.0124 | < 0.001 | Yes |
| C_xU_ vs U_xU_ | 9.6993 | < 0.001 | Yes |
| U_V_ vs U_xC_ | 3.6432 | < 0.001 | Yes |
| U_V_ vs U_xCW_ | 1.7994 | < 0.001 | Yes |
| U_V_ vs U_xU_ | 3.4862 | < 0.001 | Yes |
| U_xC_ vs U_xCW_ | -1.8439 | < 0.001 | Yes |
| U_xC_ vs U_xU_ | -0.1570 | 0.9999 | No |
| U_xCW_ vs U_xU_ | 1.6869 | < 0.001 | Yes |

**Table S5.** Tukey post-hoc comparisons of DFA axis 2 scores across all ten female treatments.

| **Comparison** | **Mean difference** | **Adjusted *p*** | **Significant** |
| --- | --- | --- | --- |
| CW_V_ vs CW_xCW_ | -0.7945 | 0.1218 | No |
| CW_V_ vs C_V_ | 0.5638 | 0.5767 | No |
| CW_V_ vs C_xC_ | 6.0553 | < 0.001 | Yes |
| CW_V_ vs C_xCW_ | -4.1265 | < 0.001 | Yes |
| CW_V_ vs C_xU_ | 3.3314 | < 0.001 | Yes |
| CW_V_ vs U_V_ | -0.3454 | 0.9641 | No |
| CW_V_ vs U_xC_ | 4.1735 | < 0.001 | Yes |
| CW_V_ vs U_xCW_ | -0.7336 | 0.2031 | No |
| CW_V_ vs U_xU_ | 3.5727 | < 0.001 | Yes |
| CW_xCW_ vs C_V_ | 1.3583 | < 0.001 | Yes |
| CW_xCW_ vs C_xC_ | 6.8498 | < 0.001 | Yes |
| CW_xCW_ vs C_xCW_ | -3.3321 | < 0.001 | Yes |
| CW_xCW_ vs C_xU_ | 4.1258 | < 0.001 | Yes |
| CW_xCW_ vs U_V_ | 0.4491 | 0.8376 | No |
| CW_xCW_ vs U_xC_ | 4.9680 | < 0.001 | Yes |
| CW_xCW_ vs U_xCW_ | 0.0609 | 1.0000 | No |
| CW_xCW_ vs U_xU_ | 4.3672 | < 0.001 | Yes |
| C_V_ vs C_xC_ | 5.4915 | < 0.001 | Yes |
| C_V_ vs C_xCW_ | -4.6904 | < 0.001 | Yes |
| C_V_ vs C_xU_ | 2.7675 | < 0.001 | Yes |
| C_V_ vs U_V_ | -0.9092 | 0.0390 | Yes |
| C_V_ vs U_xC_ | 3.6097 | < 0.001 | Yes |
| C_V_ vs U_xCW_ | -1.2974 | < 0.001 | Yes |
| C_V_ vs U_xU_ | 3.0089 | < 0.001 | Yes |
| C_xC_ vs C_xCW_ | -10.1818 | < 0.001 | Yes |
| C_xC_ vs C_xU_ | -2.7240 | < 0.001 | Yes |
| C_xC_ vs U_V_ | -6.4007 | < 0.001 | Yes |
| C_xC_ vs U_xC_ | -1.8818 | < 0.001 | Yes |
| C_xC_ vs U_xCW_ | -6.7889 | < 0.001 | Yes |
| C_xC_ vs U_xU_ | -2.4826 | < 0.001 | Yes |
| C_xCW_ vs C_xU_ | 7.4579 | < 0.001 | Yes |
| C_xCW_ vs U_V_ | 3.7812 | < 0.001 | Yes |
| C_xCW_ vs U_xC_ | 8.3001 | < 0.001 | Yes |
| C_xCW_ vs U_xCW_ | 3.3930 | < 0.001 | Yes |
| C_xCW_ vs U_xU_ | 7.6993 | < 0.001 | Yes |
| C_xU_ vs U_V_ | -3.6767 | < 0.001 | Yes |
| C_xU_ vs U_xC_ | 0.8422 | 0.1345 | No |
| C_xU_ vs U_xCW_ | -4.0649 | < 0.001 | Yes |
| C_xU_ vs U_xU_ | 0.2414 | 0.9981 | No |
| U_V_ vs U_xC_ | 4.5189 | < 0.001 | Yes |
| U_V_ vs U_xCW_ | -0.3882 | 0.9264 | No |
| U_V_ vs U_xU_ | 3.9181 | < 0.001 | Yes |
| U_xC_ vs U_xCW_ | -4.9071 | < 0.001 | Yes |
| U_xC_ vs U_xU_ | -0.6008 | 0.5750 | No |
| U_xCW_ vs U_xU_ | 4.3063 | < 0.001 | Yes |

**Table S6**. Tridecane detection frequencies across all ten female treatments.

| **Treatment** | **Detected (n)** | **Not detected (n)** | **Total (n)** | **Proportion detected** | **Wilson 95% CI lower** | **Wilson 95% CI upper** |
| --- | --- | --- | --- | --- | --- | --- |
| U_V_ | 0 | 26 | 26 | 0.000 | 0.000 | 0.129 |
| U_xU_ | 0 | 24 | 24 | 0.000 | 0.000 | 0.138 |
| U_xC_ | 0 | 22 | 22 | 0.000 | 0.000 | 0.149 |
| U_xCW_ | 26 | 0 | 26 | 1.000 | 0.871 | 1.000 |
| C_V_ | 0 | 26 | 26 | 0.000 | 0.000 | 0.129 |
| C_xU_ | 0 | 23 | 23 | 0.000 | 0.000 | 0.143 |
| C_xC_ | 0 | 22 | 22 | 0.000 | 0.000 | 0.149 |
| C_xCW_ | 24 | 0 | 24 | 1.000 | 0.862 | 1.000 |
| CW_V_ | 0 | 26 | 26 | 0.000 | 0.000 | 0.129 |
| CW_xCW_ | 26 | 0 | 26 | 1.000 | 0.871 | 1.000 |

**Overall test:** χ² = 245.0, df = 9, *p* < 0.001

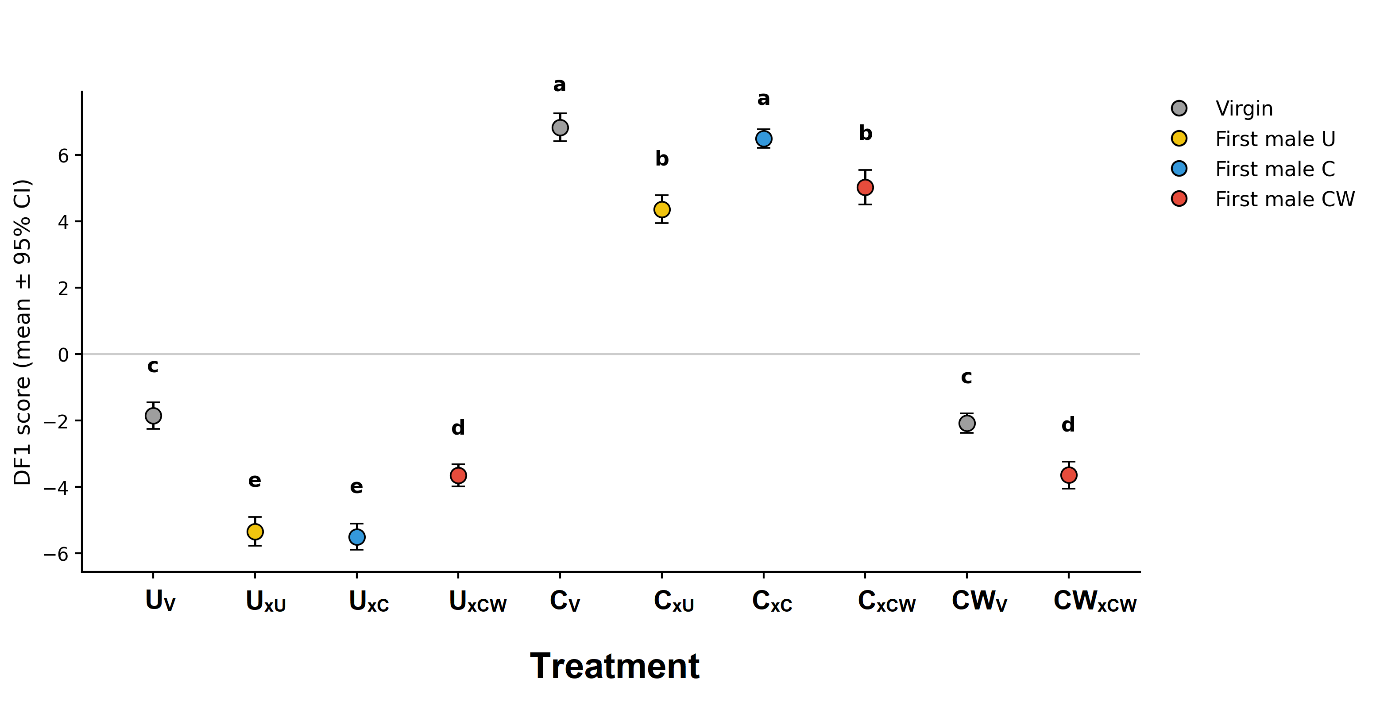

**Figure S1.** Mean DF1 scores (±95% CI) for each female treatment from the discriminant function analysis of the tridecane-excluded female CHC matrix. Different letters indicate significant differences among treatments based on Tukey post-hoc comparisons of DF1 scores.

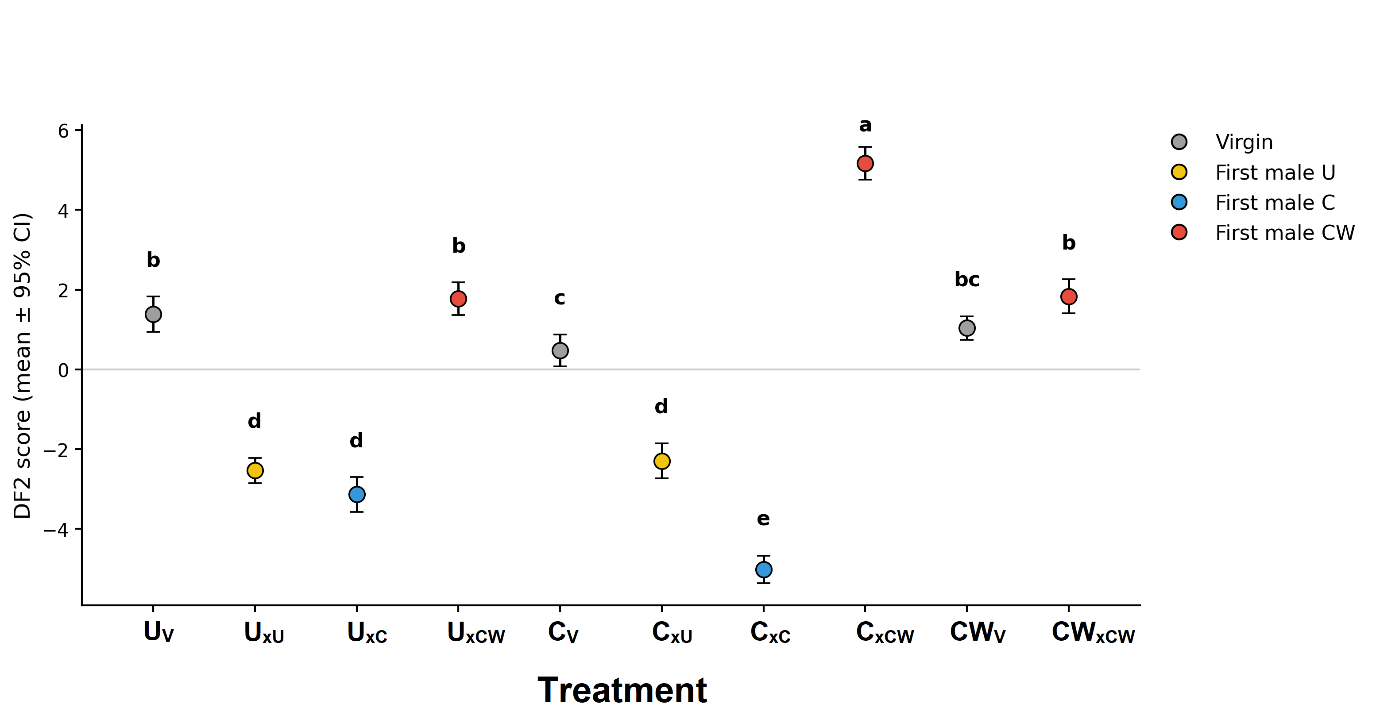
**Figure S2.** Mean DF2 scores (±95% CI) for each female treatment from the discriminant function analysis of the tridecane-excluded female CHC matrix. Different letters indicate significant differences among treatments based on Tukey post-hoc comparisons of DF2 scores.
